# Imputing abundances and inferring direction of associations between histone modifications using neural processes

**DOI:** 10.1101/2022.07.21.501033

**Authors:** Ananthakrishnan Ganesan, Denis Dermadi, Laurynas Kalesinskas, Michele Donato, Rosalie Sowers, Paul J. Utz, Purvesh Khatri

## Abstract

Histone post-translational modifications (HPTMs) play a vital role in the regulation of numerous cellular processes. However, systems level understanding of how HPTMs coordinate and interact with each other, and the direction of interactions remain unexplored due to lack of suitable technology. EpiTOF, a high-dimensional mass cytometry-based platform, measures HPTMs and histone variants across multiple immune cell sub-types at a single-cell resolution to enable the system-wide study of HPTMs. Large number of immune cells profiled using EpiTOF present an unprecedented opportunity to learn directional networks of HPTM associations. We developed a two-step computational framework to identify direction of association between two HPTMs. In the first step, we used linear regression (LR)-, *k*-nearest neighbors-(*k*NN), or Neural Processes (NP)-based to impute the abundance of HPTMs. In the second step, we developed an interpretation framework to infer direction of association between HPTMs at a system-level using mass cytometry data. We evaluated this framework using EpiTOF profiles of more than 55 million peripheral mononuclear blood cells from 158 healthy human subjects. NP models had consistently higher imputation accuracy than LR and kNN. The inferred networks recapitulated known HPTM associations and identified several novel ones. While almost all associations were conserved across all healthy individuals, in a cohort of healthy subjects, vaccinated with the trivalent inactivated seasonal influenza vaccine (TIV), we identified changes in associations between 6 pairs of HPTMs 30 days following vaccination, many of which have been shown to be functionally involved in innate memory. These results demonstrate utility of our framework in identifying causal interactions between HPTMs that can be further tested experimentally.

## Introduction

Histone post-translational modifications (HPTMs) play a vital role in the regulation of gene expression, cell differentiation, and different processes centered around DNA[1]. Dysregulation of HPTMs has been implicated in human diseases such as cancer[2, 3], infectious diseases[4], mental illnesses[5], and autoimmune disorders[6, 7]. HPTMs are also known to play an important role in the immune response following vaccination[8, 9]. Interaction of multiple HPTMs has been recognized as the language of histone crosstalk[10, 11]. Despite the advances in understanding the chromatin organization and transcriptional regulation through HPTMs, histone crosstalk has been understudied due to technological limitations of ChIP-seq[12] to investigate more than 2-3 HPTMs at once.

Recently, we described a mass cytometry-based technology, EpiTOF, for high-throughput profiling of HPTMs at a single-cell resolution[13]. EpiTOF can profile up to 38 HPTMs and histone variants across 2 non-overlapping panels. However, both panels include 11 cell phenotypic marks (CPMs) for cell type identification. Prior studies using EpiTOF profiling of immune cells have quantified increased epigenetic noise with aging in human immune system[13], identified a novel epigenetic regulation of monocyte-to-macrophage differentiation[14], and epigenetic mechanism causally linked with innate memory after vaccination[8]. Further, EpiTOF data has also elucidated comprehensive correlation networks of HPTMs that consist of modules either conserved across immune cell types or are specific to a cell lineage[15].

Despite its demonstrated advantage in identifying novel epigenetic mechanisms, EpiTOF is limited in the number of HPTMs measured in a panel due to the chemical and physical properties of heavy metal isotopes used in mass cytometry. However, the large number of cells per sample profiled using EpiTOF and recent advances in machine learning provide an opportunity to use *in silico* models to impute abundances for a subset of HPTMs, which in turn could be replaced by other HPTMs to increase the number of HPTMs profiled per cell. On the other hand, an advantage of EpiTOF is that the large number of cells profiled using it provide an unprecedented opportunity to create interpretable imputation models to infer the underlying associations between HPTMs, further elucidating the histone language[10]. Several machine learning (ML) approaches have been used to impute multi-panel cytometry data. For instance, CyTOFmerge uses a *k*-nearest neighbors (*k*NN)-based approach to create an imputation function for each subject in the dataset[16]. Such subject-specific models are advantageous since individuals are exposed to different environments that are known to have different effects on epigenetics and HPTMs[13, 17]. However, it is difficult to interpret the importance of input variables in kNN because it is non-parametric. Other approaches use parametric non-linear models such as support vector machines, boosting trees, and canonical neural networks[18]. Although these parametric models can be interpreted to quantify the importance for each input variable, they use the same imputation function for all subjects, and do not distinguish between them, which may not capture the effects of difference in environmental exposure between individuals.

Neural process (NP), a recently described neural network-based model, combines the advantages of both kNN and parametric non-linear models[19, 20]. Similar to kNN-based models, NP-based models use subject-specific functions for imputation. Because NP-based models are parametric, the importance of each input variable can be interpreted. Importantly, NP models are scalable to complex functions and large datasets[19]. Because the biological processes regulating the interactions between HPTMs are complex, we hypothesized that NPs would more accurately impute HPTMs and CPMs, and reveal biologically meaningful relationships between them.

Here, we adapted the neural network architecture from NP to impute multi-panel mass cytometry-based HPTM abundances using subject-specific parametric models. Using more than 55 million human peripheral mononuclear blood cells (PBMCs) from 158 healthy subjects across 25 independent experiments, we evaluated linear regression (LR)-, kNN- and NP-based methods for accuracy of imputation. We also developed an interpretation framework to infer direction of association between two HPTMs through systematic perturbations. We reproduced several known HPTM associations and identified several novel associations between pairs of HPTMs. Most of the associations were conserved across all healthy individuals. By combining the inferred pair-wise associations, we created a system-wide directed network of HPTM associations from the NP models. Finally, we demonstrated the utility of the NP models and the interpretation framework by identifying HPTMs whose associations were modified in response to the trivalent inactivated seasonal influenza vaccine (TIV) in healthy adults.

## Results

### Data and model types

We profiled 55.6 million PBMCs from 158 healthy subjects (13-80 years old) across 25 EpiTOF experiments and measured 38 HPTMs and histone variants, 2 core histones (H3 and H4), and 11 cell phenotypic markers (CPMs) across two panels, referred to as the methylation panel and the acetylation panel (Fig. 1A, **Fig. S1**, **Table S1**, Methods).We have previously described extensive validation of antibodies for HPTMs using Western blot, flow cytometry, and different cell lines with in vitro manipulation[13]. We divided 158 subjects into three sets: training set (15 experiments, 71 subjects, 18.9 million cells), validation set (5 experiments, 52 subjects, 20.6 million cells), and test set (5 experiments, 35 subjects, 16.1 million cells). These 25 experiments were performed over 4 years (2016-2019), collectively representing a broad range of technical heterogeneity due to different CyTOF machines, batches of reagents, and operators. We ensured that all subjects from each experiment were assigned to exactly one set for unbiased evaluation of imputation models.

**Fig. 1.**
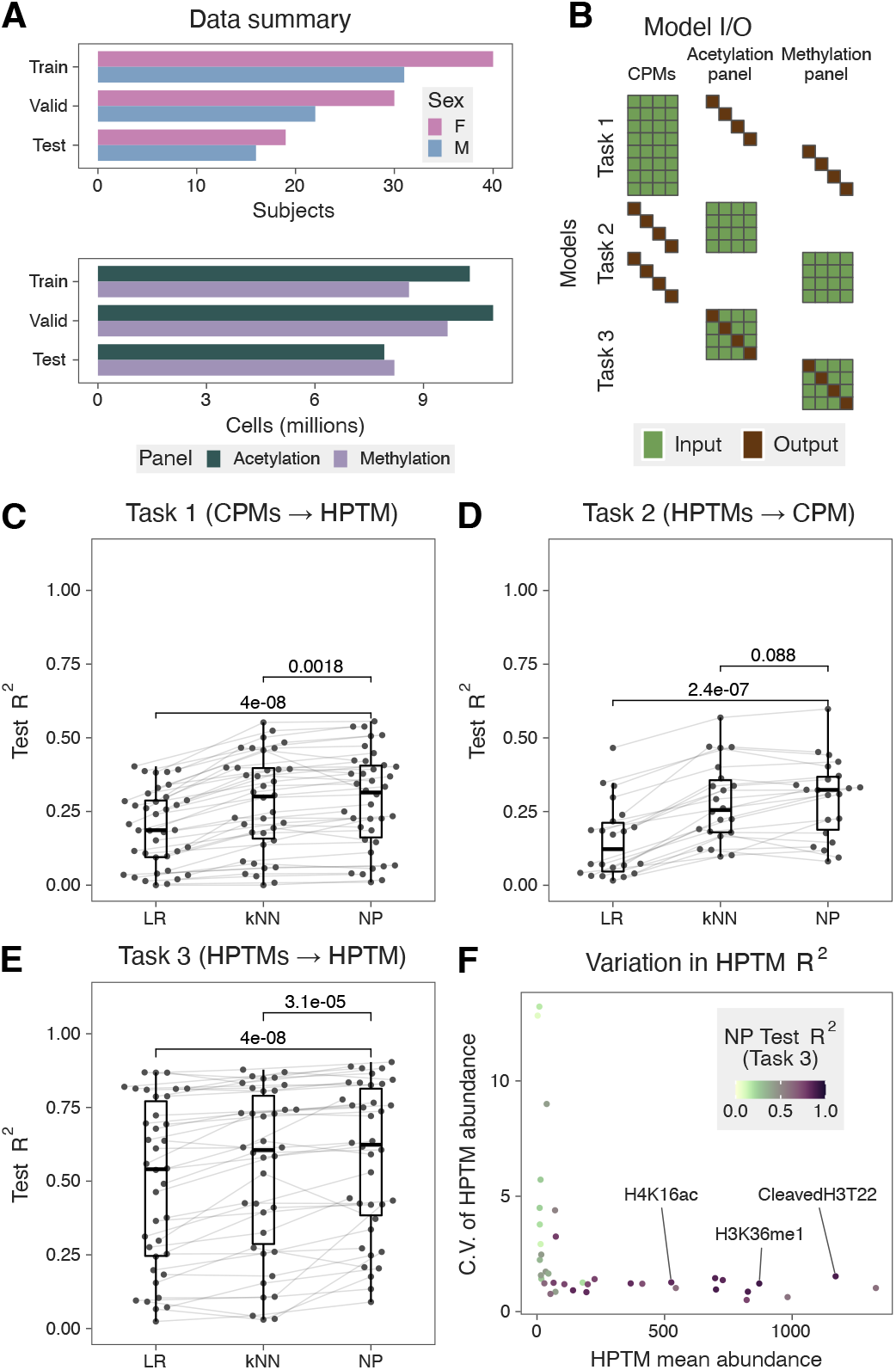
**A)** Summary of EpiTOF data. **B)** Illustration of the three model tasks and corresponding inputs and output. Each row represents a unique model, and each column represents a variable. HPTMs are split into acetylation and methylation panels. **C)** Summary of *task 1* models. Each dot represents a model. Lines connect models predicting the same HPTM across algorithms. One-sided, paired Wilcoxon test was used to compute significance of the improvement in *R*^2^ due to the NP model. **D)** Summary of *task 2* models. Each dot represents a model. Lines connect models predicting the same CPM using HPTMs from the same panel across algorithms. One-sided, paired Wilcoxon test was used to compute significance of the improvement in *R*^2^ due to the NP model. **E)** Summary of task 3 models. Each dot represents a model. Lines connect models predicting the same HPTM across algorithms. One-sided, paired Wilcoxon test was used to compute significance of the improvement in *R*^2^ due to the NP model. **F)** Comparison of *R*^2^ from task *3* NP models with mean and coefficient of variation of HPTM abundances. Each dot represents a unique HPTM. Color of each dot corresponds to the imputation *R*^2^.

CPMs are expressed on the cell surface and are used to determine the immune cell sub-type, whereas HPTMs are inside the cell nucleus. Given these biological differences between CPMs and HPTMs, we compared the accuracy of LR, *k*NN, and NP models for imputing a CPM or an HPTM using three imputation tasks: impute an HPTM using CPMs (*task 1*), impute a CPM using HPTMs (*task 2*), and impute an HPTM using other HPTMs on the same panel (*task 3*) (Fig. 1B, Methods). We used the training and validation sets for creating imputation models. Next, we locked each model and evaluated its accuracy using the test set. We evaluated 98 combinations of inputs and outputs for 294 models across the three tasks for each of the ML methods (LR, kNN, and NP). We assessed the accuracy of a model using the coefficient of determination (*R*^2^), where an *R*^2^ of 1 indicates an accurate imputation whereas an *R*^2^ of 0 indicates a failure to impute accurately. Importantly, we only report results from the test set experiments because the imputation models are expected to have high accuracy in training and validation set experiments.

### CPM and HPTM abundances are mostly independent of each other

When imputing an HPTM using CPMs in *task 1*, each of the three ML methods had low to moderate accuracy. Specifically, the mean *R*^2^ for LR, kNN, and NP models were 0.19 (range: 0-0.4), 0.28 (range: 0-0.55), and 0.29 (range: 0.01-0.56), respectively (Fig. 1C). Although NP had significantly higher mean *R*^2^ compared to LR and kNN models (*p* < 0.002), the low to moderate imputation accuracy suggests that CPMs alone are not sufficient to impute HPTMs across panels accurately.

Similarly, when imputing a CPM using HPTMs in *task 2,* each of the three methods had low to moderate accuracy. Specifically, the mean *R*^2^ for LR, kNN, and NP were 0.15 (range: 0.02-0.47), 0.28 (range: 0.10-0.57), and 0.30 (range: 0.08-0.60), respectively (Fig. 1D). NP had significantly higher mean of *R*^2^ compared to LR and kNN models (p < 0.09). Taken together, our results indicate that the abundances of CPMs and HPTMs are largely independent of each other. These results are in line with single-cell RNA-seq datasets repeatedly showing that despite the expression of the same CPM, PBMCs are highly heterogeneous[21, 22].

### HPTMs with high abundance impute each other accurately

We recently observed that HPTMs form conserved modules across immune cell types[15]. Therefore, we hypothesized that an HPTM could be more accurately imputed using other HPTMs than using CPMs. Indeed, each of the three ML models demonstrated higher accuracy for imputing an HPTM using other HPTMs (*task 3*) instead of CPMs (*task 1*) (Fig. 1E, **Fig. S2**). The mean *R*^2^ for LR, kNN, and NP were 0.49 (range: 0.02-0.87), 0.54 (range: 0.03-0.88), and 0.58 (range: 0.09-0.90), respectively. NP had significantly higher *R*^2^ compared to LR and kNN (*p* < 4e — 05). Interestingly, HPTMs imputed with high *R*^2^ were those with higher abundance and low variability across cells (Fig. 1F). It is important to note that the NP models are unaware of the mean abundance or variability of these HPTMs because the abundances of each HPTM are independently scaled by subject before being used in the models. Taken together, our results demonstrate HPTMs with high abundances are highly predictive of other HPTMs, and strongly associated with each other.

### Interpretation of NP models reveals HPTM association networks

The higher accuracy of kNN and NP in imputing an HPTM from other HPTMs (*task 3*) compared to LR suggested non-linear associations between HPTMs, which are modelled better with kNN and NP than LR. To infer pairwise HPTM associations and their directionality, we developed an interpretation algorithm using noise perturbations (Methods). Briefly, without re-training or modifying a kNN or NP model, we imputed an HPTM *Y* by systematically replacing the abundance of each input HPTM *X_i_*, one at a time, with noise. We quantified the strength of the directed association from *X_i_* to *Y* as the percent decrease in *R*^2^, with 100% denoting the strongest association and 0% denoting the weakest association. The pairs of HPTMs with strong associations formed directed networks for each panel. Interpretation of NP models identified several known directed interactions between HPTMs in both panels. In the acetylation panel, the strongest directional interaction was between histone H3 cleaved at threonine 22 (cleaved H3T22) and H3.3S31ph (Fig. 2A, **Fig. S3**). Replacing cleaved H3T22 with noise led to 90% reduction in *R* of H3.3S31ph. Importantly, perturbing H3.3S31ph had a lower effect on cleaved H3T22, with *R*^2^ reduced by 55%. The directionality of the identified association suggests biologically relevant causality, which is in line with our recent results showing that the abundance of H3.3S31ph is impacted by cleaved H3T22 in a cell line genetically modified to prevent cleavage of H3T22[14]. We also found that perturbing H3K9ac abundance reduced the accuracy of imputing PADI4 abundance, suggesting functional interaction between the two, which is in line with results from Kolodziej *et al*. demonstrating a locus-specific interaction between H3K9ac and PADI4[23].

**Fig. 2.**
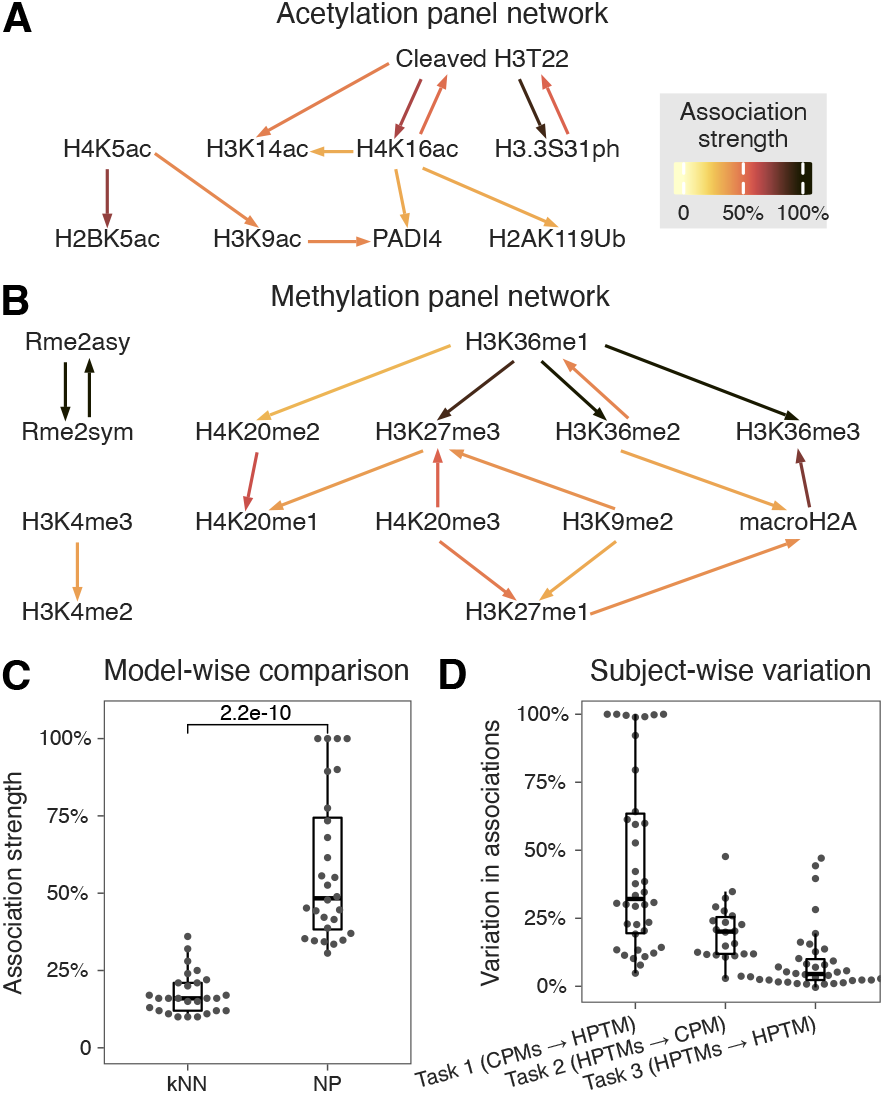
**A)** Acetylation and **B)** Methylation panel HPTM networks inferred from *task 3* (HPTMs → HPTM) NP models. Arrows go from input to output HPTM. **C)** Comparison of inferred association strengths from NP and *k*NN models. Each dot represents the association strength of a pair of HPTMs in the association networks from the corresponding models. Wilcoxon test was used to compute the significance in the difference of means. **D)** Summary of variation in model association strength across subjects. Each dot represents a model. Association strength of a model is the strength of all associations containing inputs and output of the model.

For the methylation panel, perturbation analysis identified three distinct network components (Fig. 2B, **Fig. S3**). Asymmetric and symmetric arginine dimethylation (Rme2asy/sym) strongly influenced each other and were disconnected from the lysine methylation network. This is consistent with the fact that the arginine and lysine methylations are catalyzed by different sets of enzymes. The second component consisted of H3K4me2/3, which are known to promote gene expression and occur near promoters and enhancers. The third component, on the other hand, contained known markers of gene repression and heterochromatin, including H3K27me3, H3K9me2, H4K20me3 and macroH2A. In this network component, we found that perturbing H3K9me2 abundance reduced the accuracy of imputing H3K27me3 abundance, which is in line with H3K9me2 and H3K27me3 cooperating to maintain heterochromatin[24].

In addition to these known interactions, we identified directional trans histone interactions between cleaved H3T22 and H4K16ac, and between H3K9ac, H4K5ac and H2BK5ac. We also found directional cis interactions between modifications of the same amino acid such as H3K36me1/2/3 and H4K20me2/3 (Discussion). For several HPTMs imputed with *R*^2^ > 0.5 (e.g., H3K36me2/3, Rme2asy/sym, cleaved H3T22, H3.3S31ph, H4K16ac), there was at least one input HPTM leading to > 50% decrease in *R*^2^ when its abundance was replaced with noise, suggesting highly conserved interactions between these HPTMs (**Fig. S3**). Conversely, several HPTMs imputed with *R*^2^ > 0.5 (e.g., H3K36me1, H4K20me2/3, H3K9ac, H4K5ac, PADI4, macroH2A) had no such inputs. Since the latter HPTMs were accurately imputed, detected associations suggest interactions between their inputs (Discussion). Taken together, our analyses identified several known associations between pairs of HPTMs together with their directionality. We also identified several novel associations that can be used to generate new hypotheses to further explore the crosstalk of HPTMs.

In contrast, when we applied our perturbation algorithm to the kNN models, the strength of associations was significantly lower (p = 2.2e — 10; Fig. 2C, **Fig. S4 & S5**). This is expected because kNN models are non-parametric and perturbation to any input HPTM leads to recalibration of the kNN model, which in turn can result in lower strength of associations. In contrast, NP models do not recalibrate during the perturbation analysis. Furthermore, kNN models only identified a subset of known HPTM associations (e.g., H3K4me2-H3K4me3 and Rme2sym-Rme2asy), but failed to identify the directionality of the other known associations. For instance, kNN found equally strong bidirectional association between cleaved H3T22 and H3.3S31ph (**Fig. S5**) and did not identify the association between PADI4 and H3K9ac. Overall, NP more accurately imputed HPTM abundances and better identified direction of interaction between HPTMs than kNN.

### HPTM associations are conserved across subjects

Each NP model is subject-specific and learns a unique embedding for each subject from the input data. However, we averaged the inferred HPTM associations across all subjects in the test set. Thus, it is possible that our overall associations are driven by a subset of subjects. Therefore, we modified our perturbation algorithm to quantify the between-subject variability in the associations (Methods). Briefly, we replaced the embedding for a subject with that of a different subject and estimated the between-subject variability as the percentage change in *R*^2^. Variability of 0% implies the model is independent of the subject embedding and the pair-wise associations for a given NP model are conserved, whereas variability of 100% implies the associations are distinct for each subject.

The NP models predicting an HPTM from CPMs (*task 1*) showed the highest between-subject variability (median=32.1%; Fig. 2D). This is in line with single-cell RNA-seq studies repeatedly showing large amount of heterogeneity in immune cells identified using CPMs[21, 22]. The NP models predicting a CPM from HPTMs (*task 2*) had moderate between-subject variability (median=20.1%), which is also expected since we have observed immune cell type-specific HPTM profiles that are affected by aging[13, 14]. The NP models predicting an HPTM from other HPTMs (*task 3*) had the lowest between-subject variability (median=4.5%; **Fig. S6**). Overall, our results show that the associations between CPMs and HPTMs vary between subjects, presumably affected by their different environmental exposures. On the other hand, low between-subject variability in the interactions between HPTMs suggests that most of the interactions between HPTMs are highly conserved such that changes in one HPTM lead to predictable changes in the other HPTMs.

### A subset of HPTMs accurately imputed several HPTMs

Based on low between-subject variability in predicting an HPTM using other HPTMs, combined with our previous observation of highly conserved HPTM correlation modules[15], we hypothesized that a subset of HPTMs may be more informative than others. To test this hypothesis, we ranked HPTMs using their average association strength, defined as the mean strength of all the associations containing the given HPTM as the input (Fig. 3A-B & **Fig. S3**). Cleaved H3T22 and H4K16ac in the acetylation panel, and H3K36me1 in the methylation panel were the strongest predictors of 12 other HPTMs (mean association strength > 25%), suggesting an important role for them in HPTM networks. While cleaved H3T22 and H3K36me1 were among the top four most abundant HPTMs, H4K16ac was not among the top 10 most abundant HPTMs (Fig. 1F). Hence, our results suggest that abundance of HPTM is not confounding the average strength of HPTMs associations.

**Fig. 3.**
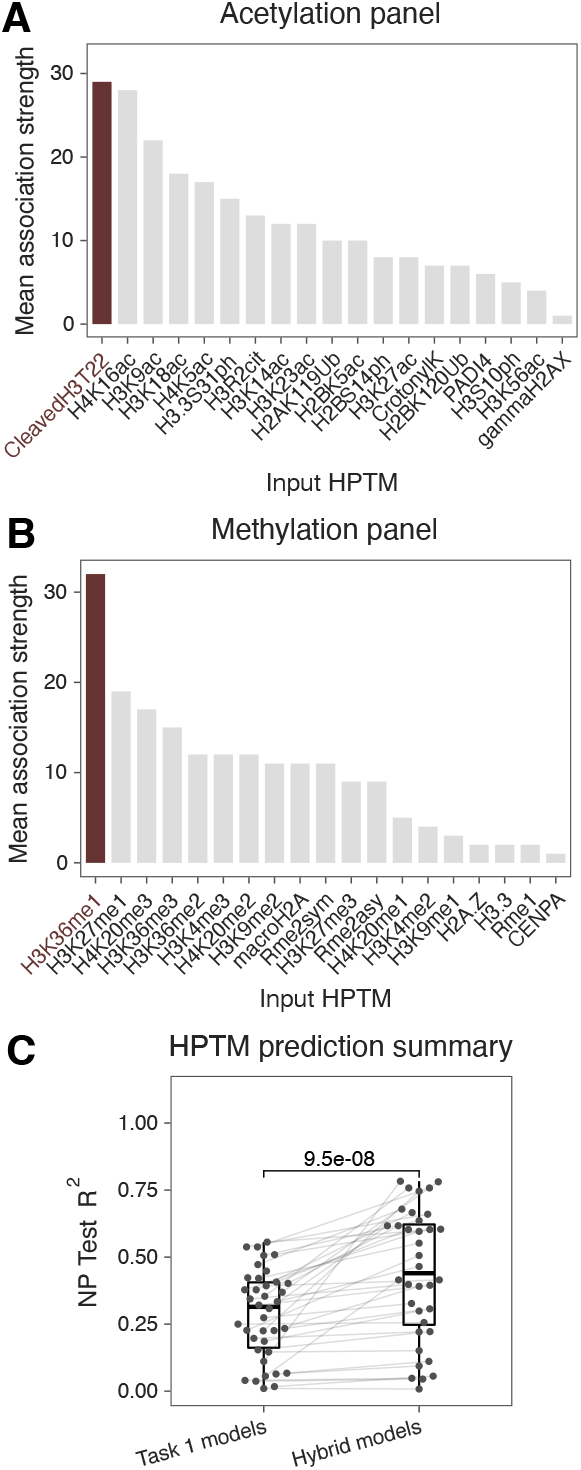
Average association strength for HPTMs in the **A)** acetylation and **B)** methylation panels. **C)** Summary of hybrid NP models that use CPMs and cleaved H3T22 (acetylation panel) or H3K36me1 (methylation panel) as inputs to impute an HPTM and comparison with models in task 1 (CPMs → HPTM). Each dot represents a unique model. Lines connect models predicting the same HPTM. One-sided, paired Wilcoxon test was used to compute significance of the improvement in *R*^2^ due to the hybrid models.

Next, we investigated whether integration of these HPTMs with CPMs would improve imputation accuracy of other HPTMs compared to using only CPMs as in *task 1.* The new hybrid NP models using 11 CPMs and either cleaved H3T22 or H3K36me1 to impute HPTMs had significantly higher *R*^2^ (mean=0.44, range: 0.01-0.78) than the models using only CPMs (p = 9.5e — 08; Fig. 3C). Importantly, the hybrid NP models had 17 HPTMs with R^2^ ≥ 0.5; more than threefold increase in accurately imputed HPTMs comparing to HPTMs imputed from only CPMs (*task 1*). Although including H4K16ac with cleaved H3T22 and 11 CPMs (mean=0.453, range: 0-0.78) significantly increased the *R*^2^ (p = 0.011) compared to including only cleaved H3T22, it did not increase the number of HPTMs imputed with *R*^2^ ≤ 0.5 (**Fig. S7**). These results show that a modified panel with H3K36me1 and cleaved H3T22 measured in both panels would enable imputing multiple HPTMs across panels with higher accuracy, further increasing the power of EpiTOF. This also highlights the power of our interpretation algorithm in accurately identifying associations and ranking the predictive power of each HPTM.

### NP models and interpretation algorithm identify influenza vaccine-associated epigenetic changes

We demonstrated that the NP models can accurately impute HPTM abundances and infer known and novel associations for healthy subjects. Next, we investigated if the same models, without any modification or re-training, could be used to impute HPTM abundances from healthy subjects affected by external stimuli such as vaccinations. We recently showed that for healthy subjects receiving the trivalent inactivated seasonal influenza vaccine (TIV), HPTM abundances are most changed 30 days after vaccination in myeloid cells, which are associated with innate memory[8]. Therefore, we evaluated the NP models imputing an HPTM from other HPTMs (*task 3*) and inferred pair-wise HPTM associations in a cohort of 21 healthy subjects sampled before (*Day 0*) and 30 days after (*Day 30*) receiving TIV (Fig. 4A, Methods). The imputation *R*^2^ at both these time points were strongly correlated with those from the healthy subjects in the test set (Fig. 4B, *R* = 0.9 and 0.87, and *p* < 1e — 12 for *Day 0* and *Day 30* samples, respectively), indicating that the same models can be used to impute HPTMs without significant change in imputation performance (**Fig. S8**). Although the inferred association strengths at both these time points were also highly correlated with those from the healthy subjects in the test set (Fig. 4C, R = 0.94 and 0.91 and *p* < 2.2*e* — 16 for *Day 0* and *Day 30* samples, respectively), some association strengths showed large deviations from the test set (**Fig. S8**), suggesting that most HPTM associations are conserved but a few are significantly modified due to TIV.

**Fig. 4.**
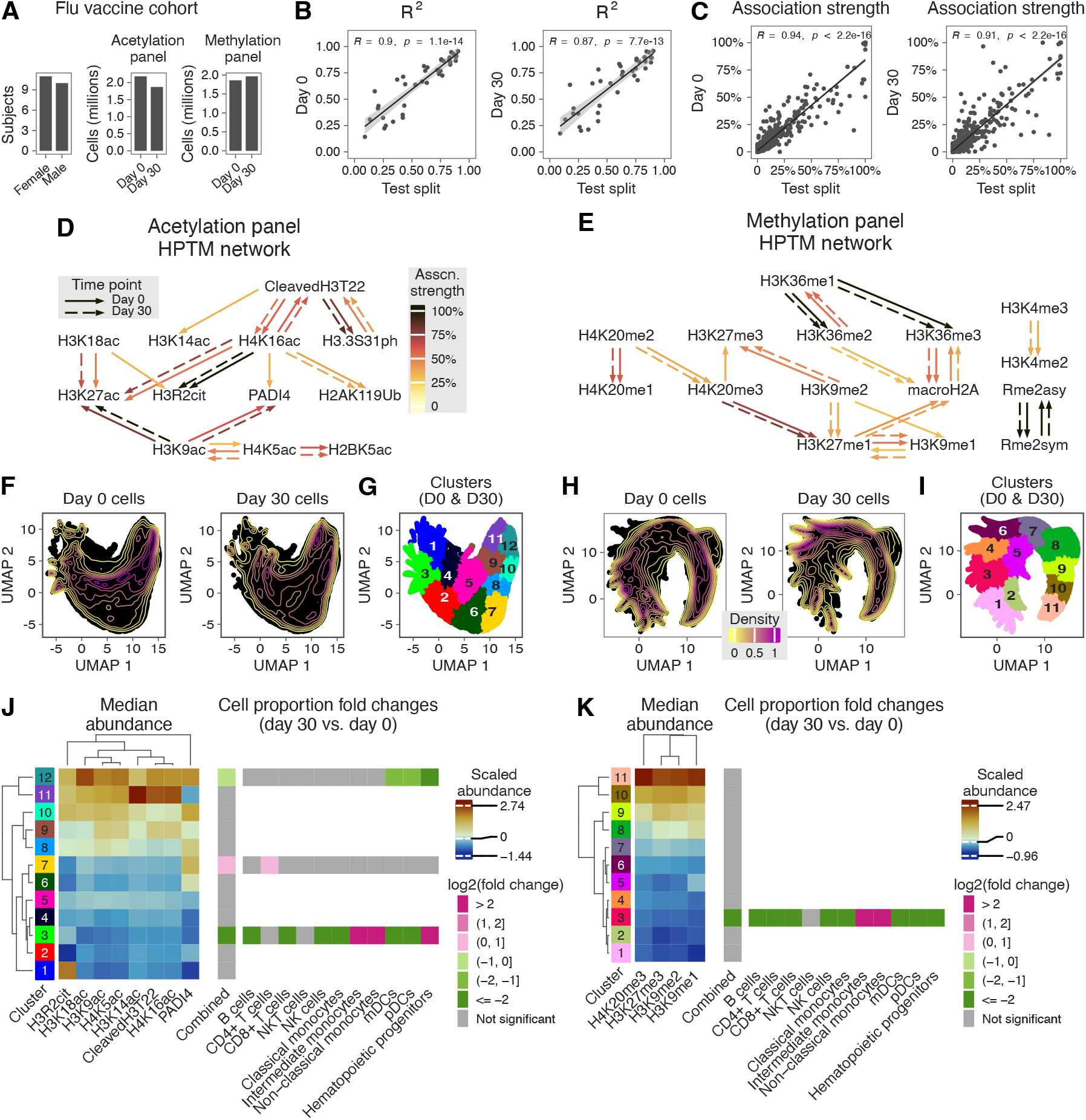
**A)** Summary of Flu vaccine cohort **B)** Comparison of NP *task 3 R^2^* between Flu vaccine cohort samples and test set. Each dot represents a unique HPTM. **C)** Comparison of HPTM association strength between Flu vaccine cohort samples and test set. Each dot represents a unique pair of HPTMs. **D)** Acetylation and **E)** methylation panel HPTM networks from *Day 0* and *Day 30* time points. Arrows go from input to output HPTM. **F)** UMAP visualization of cells in acetylation panel based on HPTMs that change in the association network (cleaved H3T22, H3K14ac, H4K16ac, PADI4, H3K18ac, H3R2cit, H2K9ac, and H4K5ac). Each time point contains 250,000 cells sampled randomly. **G)** Phenograph clustering of cells in acetylation panel. Clustering was performed in the UMAP space and used 25,000 cells per time point randomly sampled from those used for UMAP. **H)** UMAP visualization of cells in methylation panel based on HPTMs that change in the association network (H4K20me3, H3K27me3, H3K9me2, and H3K9me1). Each time point contains 250,000 cells sampled randomly. **I)** Phenograph clustering of cells in methylation panel. Clustering was performed in the UMAP space and used 25,000 cells per time point randomly sampled from those used for UMAP. Median abundance of HPTMs and fold changes in sample-wise cell proportion between *Day 30* and *Day 0* in the **J)** acetylation and **K)** methylation panels. All cells from all samples were used to calculate the medians and proportions. FDR-adjusted Wilcoxon test was used to compute significance of differences in proportion means across groups.

To identify associations that are modified due to TIV, we compared the HPTM association networks independently inferred from the *Day 0* and *Day 30* samples. 12 out of 16 *Day 0* associations in the acetylation panel network and 17 out of 19 *Day 0* associations in the methylation panel network were conserved with identical association strengths in *Day 30* samples (Fig. 4D-E). However, in the acetylation panel, association between cleaved H3T22 and H3K14ac, H4K16ac and PADI4, H3K18ac and H3R2cit, and H2K9ac and H4K5ac were present only on *Day 0* but not on *Day 30* (Fig. 4D). In the methylation panel, association between H4K20me3 and H3K27me3, and H3K9me2 and H3K9me1 were present only on *Day 0* but not on *Day 30* (Fig. 4E). Across both panels, all the HPTM pairs associated on *Day 30* were also associated on *Day 0*. Taken together, the networks revealed that the associations between 6 pairs of 12 HPTMs were significantly different on *Day 30* post-vaccination.

Furthermore, across both EpiTOF panels, PBMCs from the two time points were distributed differently in the UMAP[25] space defined by these HPTMs (Fig. 4F & 4H). To further quantify these distinct patterns, we clustered[26] the PBMCs (Fig. 4G & 4I), computed the median abundance of the HPTMs for each cluster in each panel, and compared the sample-wise proportion of cells in a cluster across the two time points ((Fig. 4J-K, Methods). In the acetylation panel, the cell proportions were significantly different in clusters *3 (FDR* = 7.2%), *7 (FDR* = 4.1%), and *12 (FDR* = 5.1%) (Fig. 4J & **Fig. S9**). Similarly, in the methylation panel, the cell proportions were significantly different in cluster *3 (FDR* = 8.4%) (Fig. 4K & **Fig. S9**). These suggest that the changes in these HPTM associations are driven by change in cell proportions in these clusters.

We have previously shown that the overall proportions of immune cell sub-types are not significantly different between *Day 30* and *Day 0* samples[8]. However, based on the clustering analysis, we hypothesized that the proportions of immune cell sub-types could be significantly different in the clusters where the overall cell proportions are significantly different. Therefore, we compared the cell proportions within clusters 3, 7, and *12* in the acetylation panel and cluster *3* in the methylation panel (Fig. 4J-K & **Fig. S9**). Cluster *3* in the acetylation panel was defined by low abundance of the HPTMs and was dominated by cells from *Day 0* samples. Importantly, in this cluster, the proportions of B cells, CD8+ T cells, NK cells, classical monocytes, mDCs, and pDCs significantly decreased, whereas those of intermediate and non-classical monocytes, and hematopoietic progenitors significantly increased in *Day 30* samples compared to *Day 0*. The proportions of intermediate and non-classical monocytes are known to increase in response to infections and inflammatory conditions[27]. In contrast, cluster *7*, which was also defined by low abundances of several HPTMs and moderate abundance of PADI4, had significantly higher proportion of cells from *Day 30* samples, which was solely due to a significant increase in CD4+ T cell proportions. Cluster 12, defined by moderate to high abundances of the HPTMs, had significantly lower proportion of cells from *Day 30* samples compared to *Day 0*, which was due to significantly reduced proportions of mDCs, pDCs, and hematopoietic progenitors on *Day 30* post-vaccination. In the methylation panel, the proportion of cells from *Day 30* samples were significantly decreased in cluster *3* compared to *Day 0*. In this cluster, defined by low abundance of the HPTMs, the proportions of B cells, CD4+ and CD8+ T cells, NK cells, classical monocytes, mDCs, pDCs, and hematopoietic progenitors decreased significantly in *Day 30* samples compared to *Day 0.* However, the proportions of intermediate and non-classical monocytes were significantly higher, similar to cluster *3* in acetylation panel. Thus, NP models and perturbation analysis led to the identification of HPTMs whose associations changed due to TIV, and clusters defined by these HPTMs showed significant differences in cell proportions. The observation of cluster-specific cell proportion changes, but not at the overall level, suggests epigenetic reprogramming of the immune cells in these clusters, which is in line with recent studies demonstrating that memory in innate immune system is driven by epigenetic reprogramming[9, 28].

In summary, we have shown that the NP models trained using EpiTOF profiles of healthy subjects can be used to impute HPTM abundances in post-vaccinated subjects without any change in imputation accuracy. The associations between 6 pairs of HPTMs are modified due to TIV, and the clusters of cells that possibly drive these differences in associations show differences in proportions of immune cell sub-types.

## Discussion

We profiled 38 HPTMs and histone variants with 11 CPMs across 2 panels in more than 55 million PBMCs from 158 healthy controls across 25 experiments using EpiTOF. Using this unique dataset, we developed a computational framework for inferring directional associations between two HPTMs. As part of the development, we evaluated three machine learning methods for imputation to conclusively show that NP models impute HPTMs and CPMs with significantly higher accuracy, measured as *R*^2^, compared to LR and *k*NN models.

Increased accuracy of NP models required larger amounts of training data, computational resources, and computational time to train than LR and *k*NN models. However, once trained, NP models offered several advantages over LR and *k*NN models. First, NP models better captured associations from the underlying data than *k*NN models and provided novel biological insights. Second, NP models were faster than *k*NN models when making predictions during testing. Third, the learnings from the trained NP models can be transferred to other settings using transfer learning approaches. For instance, when a new HPTM is added to the EpiTOF panel, the data and computational time required for training can be significantly reduced by using transfer learning approaches. Importantly, the NP approach can be applied to impute other multi-panel data sources such as CyTOF. Taken together, the advantages of the NP models outweigh its limitations for increased resources.

We found that the CPMs and HPTMs impute each other with low to moderate *R*^2^. The best imputed CPM (using HPTMs as input) was CD11c (*R*^2^ = 0.6), which is abundant only in cells from the myeloid lineage. This suggests a distinct pattern of HPTM abundances in the myeloid and lymphoid lineages. Indeed, we recently found that myeloid and lymphoid cells follow epigenetically distinct trajectories during their differentiation from hematopoietic progenitors[15]. However, other CPMs were imputed with *R*^2^ < 0.5. This is not surprising, since these CPMs delineate PBMCs into broad immune cell subtypes (e.g., CD4+/CD8+ T cells, classical/non-classical monocytes) whereas chromatin states, and hence HPTM abundances, which lead to mRNA expression, show substantial variability within the same cellular population[29].

We found that the HPTMs accurately imputed other HPTMs, and the perturbation-based analysis allowed inferring directionality between a pair of HPTMs. Our analysis correctly identified several known HPTM interactions. Importantly, our analyses also identified several novel associations and HPTM networks that should be further investigated in the future studies to decode the crosstalk between HPTMs. For example, we identified directed interactions between mono-, di- and tri-methylation of lysine 36 on histone H3. Although H3K36me2 and H3K36me3 have been well studied, not much is known about H3K36me1[30]. *NSD1* is known to catalyze H3K36me1/2 and *SETD2* catalyzes H3K36me3 *in vivo.* However, it is unclear whether the substrate for the *SETD2*-dependent catalysis of H3K36me3 is H3K36me1, H3K36me2, or both[31]. Our perturbation analysis strongly suggests that H3K36me1, not H3K36me2, is a stronger predictor of H3K36me3, which in turn suggests that H3K36me1 is the substrate for *S’ETD2*-dependent catalysis of H3K36me3.

We identified cleaved H3T22 and H3K36me1 as strong predictors for HPTMs in the acetylation and methylation panels respectively and showed that a hybrid model using cleaved H3T22 and H3K36me1 with the CPMs significantly improves imputation accuracy for HPTMs. We note that both cleaved H3T22 and H3K36me1 are not measured in ENCODE[32] and are understudied[30, 33]. Our results indicate a pair of highly abundant yet understudied HPTMs to play a central role in HPTM associations and crosstalk.

The inferred HPTM associations also revealed several accurately imputed HPTMs (e.g., H3K36me1, H4K20me2/3, H3K9ac, H4K5ac, PADI4, and macroH2A) that had no single strong predictor. These likely suggest that several predictors interact with each other such that the removal of a single predictor doesn’t significantly affect the imputation of these HPTMs. Inferring these interactions would require performing perturbation analyses on several combinations of inputs for a single output. In this work, we only focus on pair-wise associations between an input and an output. Second, we did not analyze each immune cell sub-type separately in this proof-of-concept study. Hence, our analysis does not identify the immune cell sub-type where these associations are identified. As more data becomes available, future studies should train NP models per cell type to potentially further increase the accuracy of imputation and to derive cell type-specific HPTM associations.

In summary, we described a machine learning-based framework to impute abundances for a subset of HPTMs with high accuracy. We found that NP-based models, which had the highest accuracy, have several advantages compared to other imputation methods. We also demonstrated a novel interpretation algorithm that infers valuable insights into the underlying biological processes from the data. Our methods have potential to be applied to other single-cell frameworks like CyTOF and scRNA-seq to infer directed associations.

## Methods

### EpiTOF data

The 158 healthy subjects were enlisted through 6 cohorts and their EpiTOF profiles were measured across 25 experiments (**Table S2**). We have previously described all 6 cohorts-the Atlanta cohort[8] (50 subjects, 13 experiments), BR (24 subjects, 2 experiments) and Twins cohorts[13] (40 subjects, 2 experiments), Oklahoma cohort[34] (18 subjects, 4 experiments), Stanford (16 subjects, 3 experiments) and South Africa cohorts[15] (10 subjects, 1 experiment). The effects of TIV were studied on 21 subjects from the Atlanta cohort. The *Day 30* time point samples were not present in any of training, validation, or test sets.

### Imputation tasks, model inputs and outputs

We evaluated models imputing a CPM or an HPTM across 4 tasks: an HPTM imputed using CPMs (*task 1*), a CPM imputed using HPTMs (*task 2*), an HPTM imputed using other HPTMs on the same panel (*task 3*), and an HPTM imputed using CPMs and best predicting HPTM (hybrid models). Each model also uses the abundances of H3 and H4, age, and sex of the subject as inputs.

### Data scaling and splitting

We scaled each CPM and HPTM by subject to have mean 0 and standard deviation 1. Age was divided by 100 and sex was one-hot encoded. We split each experiment into exactly one of train, validation, or test set (**Table S2**).

### NP models: Design overview

We first describe how a trained and tested model will be used for imputing an HPTM across panel and use the example of imputing H3K36me1 (Y) from CPMs, H3, H4, age and sex (X) (*task 1*) for a subject *s.* The NP model consists of two neural networks-the *encoder* and the *imputer*. The *encoder* processes cells from the panel where *Y* is measured and known. For H3K36me1, it is the methylation panel. We denote the cells processed by the *encoder* as the *context cells*. First, the *encoder* encodes each *context cell*:

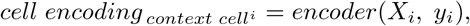

where

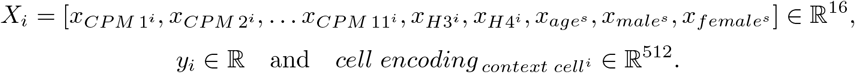

*x*_*CPM* 1^*i*^_ and *y_i_* represent the measured abundance of CPM 1 and H3K36me1 respectively in the *i^th^ context cell* for subject *s*.

Next, the subject embedding for *s* is calculated as

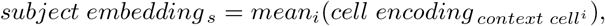

where *subject embedding_s_* ∈ ℓ^512^.

The *imputer* processes cells from the panel where Y needs to be imputed. For H3K36me1, it is the acetylation panel. We denote the cells processed by the *imputer* as the *target cells.* The *imputer* imputes H3K36me1 abundance for each *target cell*:

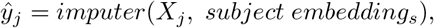

where *ŷ_j_* ∈ ℝ represents the imputed abundance of H3K36me1 in the *j^th^ target cell.*

### NP models: Training and testing

For training and testing NP models, we only used cells from the panel where Y is measured. We randomly divided the cells from this panel into two equal parts and used the first as the *context cells* and the second as the *target cells.* Due to GPU memory limitations, we split cells from each subject into 20 parts randomly and processed each as a separate subject.

We computed the training loss as the sum of two terms

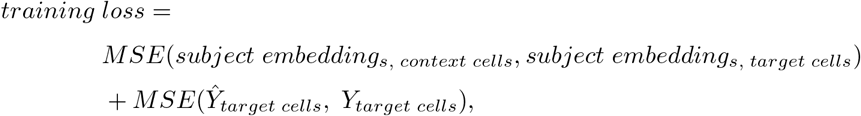

where *MSE*() is the mean squared error, *subject embedding_s, context cells_* ∈ ℝ^512^ and *subject embedding_s, target cells_* ∈ ℝ^512^ are computed by the *encoder* using the *context* and *target cells* respectively, and *Y_target cells_* ∈ ℝ^*nt*^ and *Ŷ_target cells_* ∈ ℝ^*nt*^ are the measured and the *imputer* imputed values respectively for the output corresponding to the nt *target cells.* The first term corresponds to the *encoder* network loss and the second the *imputer* network loss.

We used the coefficient of determination on the test split to assess the model performance:

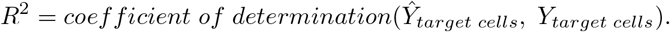

### NP models: Architecture

The *encoder* and *imputer* networks are fully connected with 2 hidden layers of dimension 256 (**Table S3**). The input dimensions for each network varies by task. The *encoder* output has dimension 512 and the *imputer* output is a scalar.

### *k*NN and LR models

*k*NN models impute Y for a *target cell* based on its nearest neighbors in the *context cells.* To be comparable with the NP models, the division of cells into *context* and *target* groups are the same for both NP and *k*NN. We evaluated *k* ∈ {1, 5,10, 20, 50,100, 200} on the validation set and used the best *k* for each model on the test set.

LR models impute Y for a *target cell* based on the learned linear combination of the inputs *X*. The LR models do not use the *context cells*.

### Perturbation-based algorithm to infer HPTM associations

To infer the association strength between an input variable *X_i_* and output *Y*, we designed a perturbation-based interpretation algorithm. First, we replaced the measured values of *X_i_* in the *target cells* for subject s with random values drawn from 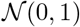 and imputed the output variable:

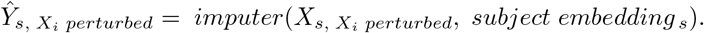

We then evaluated *R*^2^ after input perturbation:

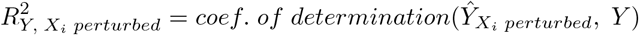

and then obtained the strength of the directed association from *X_i_* to *Y*:

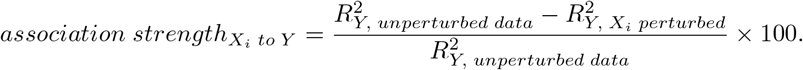

### Constructing directed HPTM association network from inferred pair-wise association strengths

To obtain the association network for HPTMs in a panel from the pair-wise association strengths, we first removed HPTMs imputed with *R*^2^ ≤ 0.5. Among the associations between the remaining HPTMs, we calculated the threshold strength as

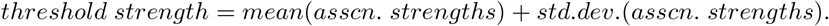

We then removed all associations with strength lesser than the threshold strength and combined the remaining associations into a single directed network.

### Perturbation-based algorithm to infer subject-wise variation in NP models

To infer the subject-wise variation in an NP model, we replaced the computed *subject embedding_s_* for subject s with those from a different random subject *ŝ* and imputed the output variable:

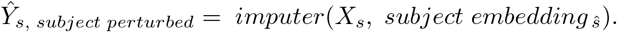

We then evaluated *R*^2^ after input perturbation:

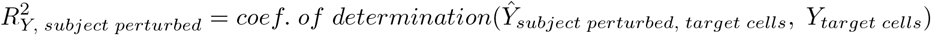

and then obtained the subject-wise variation for the NP model:

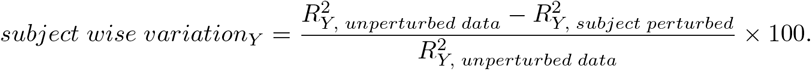

*subject wise variation_Y_* = 0% implies the model, and the underlying associations, are independent of the subject. The greater this value, the more the variation in the underlying associations across subjects.

### Dimensionality reduction and clustering for studying the effects of TIV

We computed UMAP[25] projections on a subset of 1,000,000 cells (250,000 randomly sampled cells from each panel and time point) using *n neighbors* = 15 and *min dist* = 0.1. We performed phenograph clustering[26] on the UMAP space using a subset of 100,000 cells (25,000 randomly sampled cells from those used for UMAP from each panel and time point) using *k* = 1000. We then transformed every cell from both the time points to the UMAP space without modifying the UMAP projection function and assigned clusters based on the 5 nearest neighbors in the subset where the clusters were computed. We computed the cluster-wise median HPTM abundances and sample-wise cell proportions using all the cells.

## Contributions

PK conceived the study. PK and PJU obtained funding. AG performed computational analyses and wrote code with help from RS. AG, DD, LK, and MD interpreted data. PJU supervised EpiTOF profiling. AG and PK wrote the manuscript with contributions from all coauthors. PK and PJU supervised the study. All authors approved the manuscript.

## Supporting information

Compiled supplementary figures and tables

