## Supplementary material for "Imputing abundances and inferring direction of associations between histone modifications using neural processes": Compiled supplementary figures and tables

**Table S1** CPMs and HPTMs measured in the EpiTOF data

| CPMs | HPTMs (Acetylation panel) | HPTMs (Methylation panel) |
|---|---|---|
| CD11c | CleavedH3T22 | CENPA |
| CD123 | CrotonylK | H2A.Z |
| CD14 | gammaH2AX | H3.3 |
| CD16 | H2AK119Ub | H3K27me1 |
| CD19 | H2BK120Ub | H3K27me3 |
| CD3 | H2BK5ac | H3K36me1 |
| CD4 | H2BS14ph | H3K36me2 |
| CD45 | H3.3S31ph | H3K36me3 |
| CD56 | H3K14ac | H3K4me2 |
| CD8 | H3K18ac | H3K4me3 |
| HLADR | H3K23ac | H3K9me1 |
| | H3K27ac | H3K9me2 |
| | H3K56ac | H4K20me1 |
| H3 | H3K9ac | H4K20me2 |
| H4 | H3R2cit | H4K20me3 |
| | H3S10ph | macroH2A |
| | H4K16ac | Rme1 |
| | H4K5ac | Rme2asy |
| | PADI4 | Rme2sym |

**Table S2** Summary of EpiTOF experiments.

| Set | Experiment | Subjects |
|---|---|---|
| Training | Atlanta cohort 1 | 4 |
| | Atlanta cohort 2 | 4 |
| | Atlanta cohort 3 | 4 |
| | Atlanta cohort 4 | 4 |
| | Atlanta cohort 5 | 4 |
| | Atlanta cohort 6 | 4 |
| | Atlanta cohort 7 | 3 |
| | Atlanta cohort 8 | 4 |
| | Atlanta cohort 9 | 4 |
| | Stanford cohort 1 | 6 |
| | Stanford cohort 2 | 3 |
| | Stanford cohort 3 | 7 |
| | South Africa cohort | 1 |
| | Oklahoma cohort 1 | 5 |
| | Oklahoma cohort 2 | 5 |
| Validation | Twins cohort 1 | 20 |
| | Twins cohort 2 | 20 |
| | Atlanta cohort 10 | 4 |
| | Oklahoma cohort 3 | 4 |
| | Oklahoma cohort 4 | 4 |
| Test | BR cohort 1 | 12 |
| | BR cohort 2 | 12 |
| | Atlanta cohort 11 | 4 |
| | Atlanta cohort 12 | 3 |
| | Atlanta cohort 13 | 4 |
| Flu vaccine cohort | Atlanta cohort 1 | 2 |
| | Atlanta cohort 2 | 1 |
| | Atlanta cohort 3 | 2 |
| | Atlanta cohort 4 | 4 |
| | Atlanta cohort 5 | 4 |
| | Atlanta cohort 6 | 2 |
| | Atlanta cohort 12 | 2 |
| | Atlanta cohort 13 | 4 |

**Table S3** NP architecture.

| *Encoder* dimensions | | | | |
|---|---|---|---|---|
| Task | Input | Hidden 1 | Hidden 2 | Output |
| *Task 1* (CPMs → HPTM) | 16 | 256 | 256 | 512 |
| *Task 2* (HPTMs → CPM) | 24 | 256 | 256 | 512 |
| *Task 3* (HPTMs → HPTM) | 23 | 256 | 256 | 512 |
| *Hybrid models* | 17 | 256 | 256 | 512 |
| *Imputer* dimensions | | | | |
| Task | Input | Hidden 1 | Hidden 2 | Output |
| *Task 1* (CPMs → HPTM) | 527 | 256 | 256 | 1 |
| *Task 2* (HPTMs → CPM) | 535 | 256 | 256 | 1 |
| *Task 3* (HPTMs → HPTM) | 534 | 256 | 256 | 1 |
| *Hybrid models* | 528 | 256 | 256 | 1 |

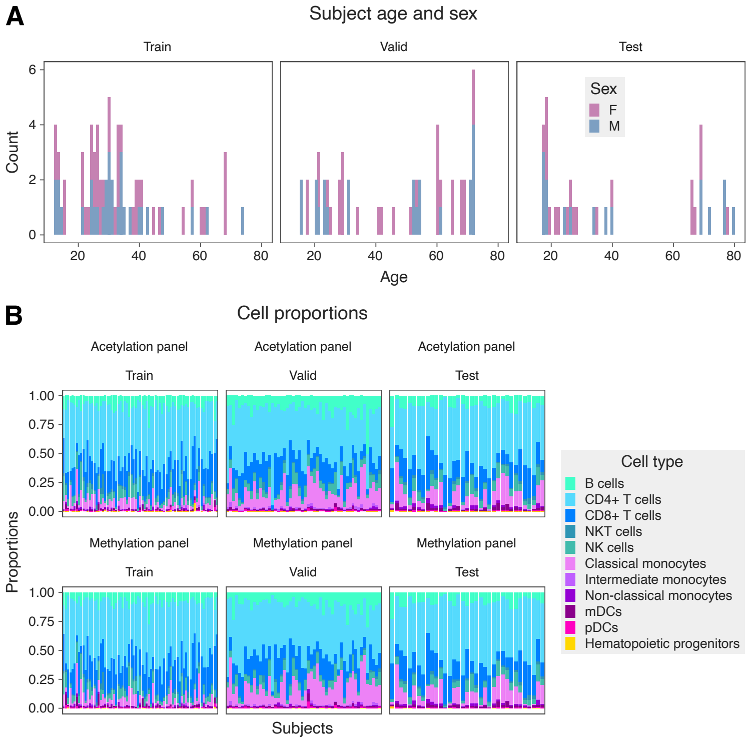


**Fig. S1** **A)** Distribution of age and sex in training, validation, and test sets. B) Distribution of cell proportions for the 11 immune cell sub-types. Each bar corresponds to a subject.

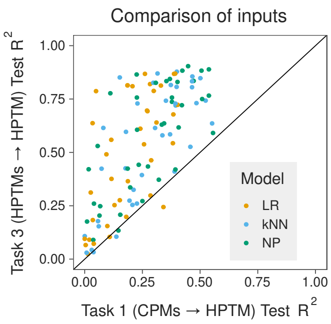


**Fig. S2** Comparison of models in tasks 1 and 3. Each dot represents a unique combination of HPTM and algorithm.

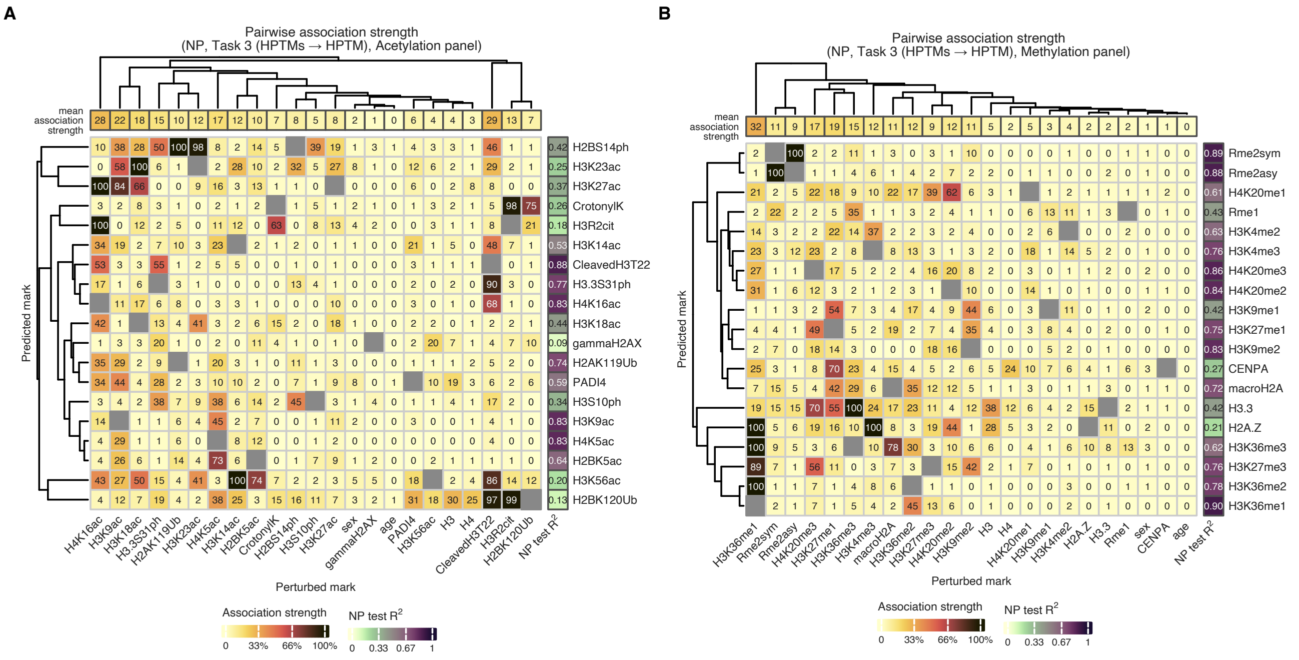


**Fig. S3** NP-inferred association strength in test set for pairs of HPTM in **A)** acetylation and **B)** methylation panel.

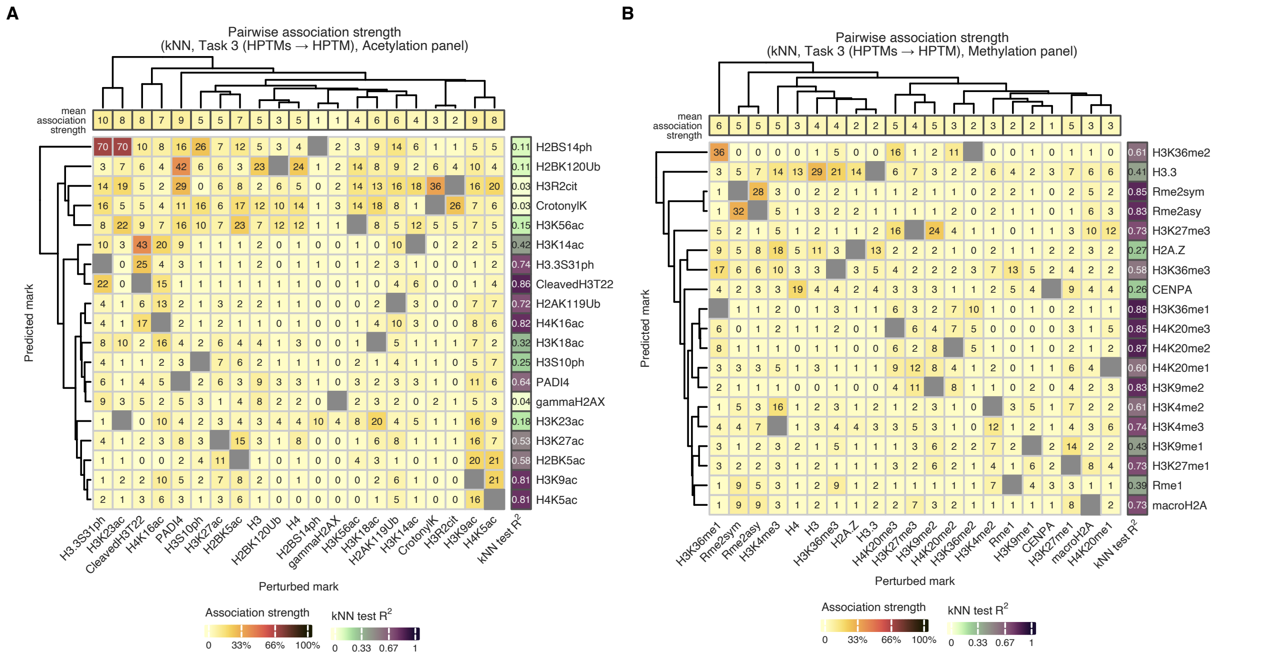


**Fig. S4** *k*NN-inferred association strength in test set for pairs of HPTM in **A)** acetylation and **B)** methylation panel.

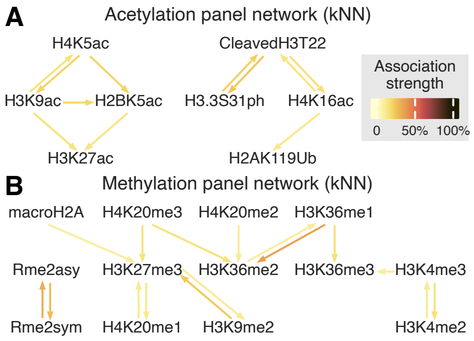


**Fig. S5** *k*NN-inferred association network in test set for HPTMs in **A)** acetylation and **B)** methylation panel.

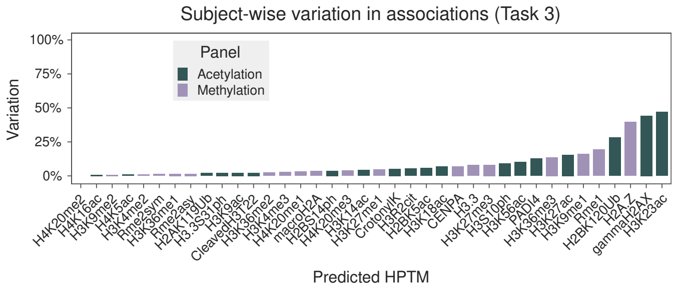


**Fig. S6** Subject-wise variation in association for each NP model in *task 3*.

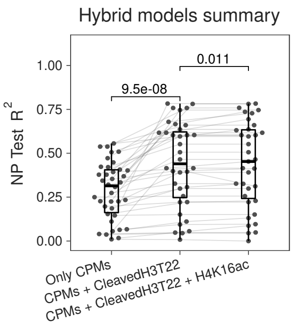


**Fig. S7** Summary of hybrid models. Each dot corresponds to an HPTM. One-sided, paired Wilcoxon test was used to compute significance of the improvement in $R^2$.

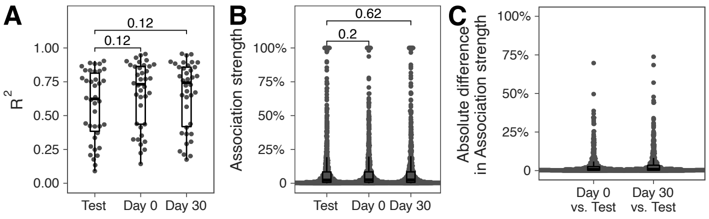


**Fig. S8** **A)** Comparison of $R^2$ between test set, *Day 0*, and *Day 30*. Each dot represents an HPTM. FDR-adjusted Wilcoxon test was used to compute significance of differences in proportion means across groups. **B)** Comparison of association strength between test set, *Day 0*, and *Day 30*. Each dot represents a unique pair of HPTMs. FDR-adjusted Wilcoxon test was used to compute significance of differences in proportion means across groups. **C)** Absolute difference in association strength between test set *vs. Day 0* and *Day 30* time points. Each dot represents a unique pair of HPTMs.

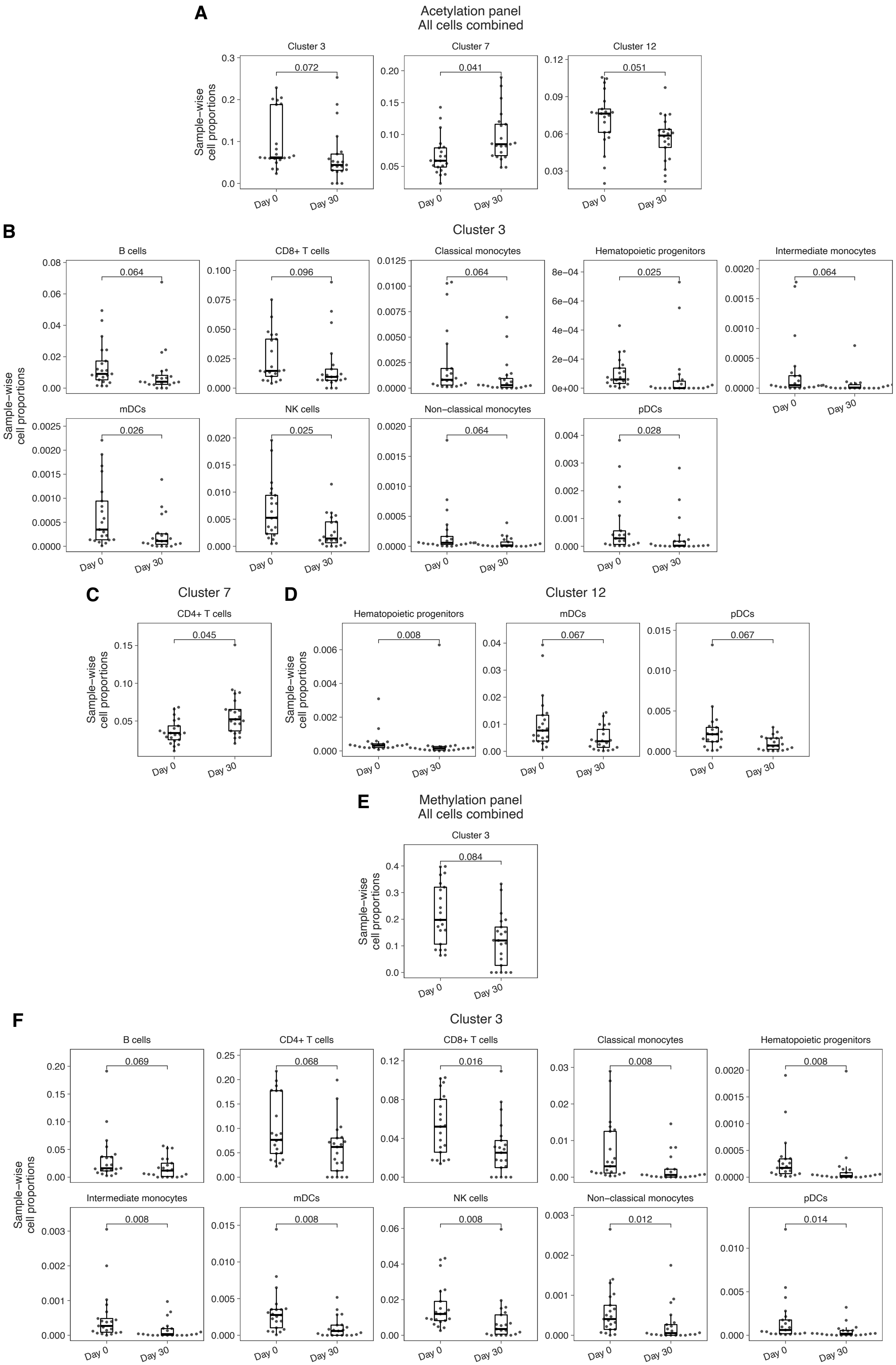


**Fig. S9** Sample-wise cell proportions in **A)** acetylation panel, all cell immune cell sub-types combined, **B)** acetylation panel, cluster *3*, by immune cell sub-type, **C)** acetylation panel, cluster *7*, by immune cell sub-type, **D)** acetylation panel, cluster *12*, by immune cell sub-type, **E)** methylation panel, all immune cell sub-types combined, and **F)** methylation panel, cluster *3*, by immune cell sub-type. Each dot represents a subject. All cells from all samples were used to calculate the proportions. FDR-adjusted Wilcoxon test was used to compute significance of differences in proportion means across groups.
